# Morphometric and Functional Brain Connectivity Differentiates Chess Masters from Amateur Players

**DOI:** 10.1101/2020.09.18.303685

**Authors:** Harish RaviPrakash, Syed Muhammad Anwar, Nadia M. Biassou, Ulas Bagci

## Abstract

A common task in brain image analysis includes diagnosis of a certain medical condition wherein groups of healthy controls and diseased subjects are analyzed and compared. On the other hand, for two groups of healthy participants with different proficiency in a certain skill, a distinctive analysis of the brain function remains a challenging problem. In this study, we develop new computational tools to explore the functional and anatomical differences that could exist between the brain of healthy individuals identified on the basis of different levels of task experience/proficiency. Towards this end, we look at a dataset of amateur and professional chess players, where we utilize resting-state functional magnetic resonance images to generate functional connectivity (FC) information. In addition, we utilize T1-weighted magnetic resonance imaging to estimate morphometric connectivity (MC) information. We combine functional and anatomical features into a new connectivity matrix, which we term as the *functional morphometric similarity connectome* (*FMSC*). Since, both the FC and MC information is susceptible to redundancy, the size of this information is reduced using statistical feature selection. We employ off-the-shelf machine learning classifier, support vector machine, for both single- and multi-modality classifications. From our experiments, we establish that the saliency and ventral attention network of the brain is functionally and anatomically different between two groups of healthy subjects (chess players). We argue that, since chess involves many aspects of higher order cognition such as systematic thinking and spatial reasoning and the identified network is task-positive to cognition tasks requiring a response, our results are valid and supporting the feasibility of the proposed computational pipeline. Moreover, we quantitatively validate an existing neuroscience hypothesis that learning a certain skill could cause a change in the brain (functional connectivity and anatomy) and this can be tested via our novel FMSC algorithm.

## Introduction

Functional connectivity networks (FCNs) are representative of relationships between spatially separated brain regions. The study of FCN, as a technique for diagnosis of clinical conditions, has gained an increase in popularity due to their high test-retest reliability and reproducibility^1^. There are computational methods proposed to generate FCNs which can delineate similarities and differences between healthy controls and diseased subjects^2–4^. While methods that use machine learning approaches for analyzing FCNs have shown great promise^5–7^, FCN-based classification methods could suffer from high dimensionality issues where there are more features than the available data. The use of feature selection algorithms with sparse learning is a common approach employed to handle such feature dimensionality problems. Sparse learning techniques, for instance, were successfully applied for diagnosis of Alzheimer’s disease (AD)^8^, attention deficit hyperactivity disorder (ADHD)^9^, and epilepsy^10^. While there is evidence that FCN-based methods are useful for studying various clinical conditions, these have not yet been widely adapted to analyze two groups of healthy subjects. Structural and functional brain studies have shown that there are structural correlates of intelligence where for *intelligent* people, the brain regions are known to be more connected^11^. We propose to extend the applicability of sparse learning methods from diseased subjects to healthy subjects consisting of professional chess players (grand masters) and amateurs.

Skill acquisition, which could include long-term practicing of a particular set of actions, can lead to changes in the brain structure. For instance, in rats trained towards a reaching task with a single fore-paw, an increase in strength of horizontal connections was observed in the motor cortex^12^. The effect of long-term skill acquisition on human subjects was studied by evaluating changes in the gray matter volume using voxel-based morphometry (VBM)^13^. In another study, VBM was used to study the morphological changes associated with complexity of navigation induced learning in the brain for London taxi and bus drivers^14^. An increase in the mid-posterior hippocampus gray matter volume was observed in taxi drivers, who tend to remember complex navigation details better, compared to bus drivers. For another example, for a group of professional badminton players, altered functional connectivity patterns were found between the left superior parietal and frontal regions when compared to controls^15^. It was hypothesized that even short-term skill acquisition, such as learning to juggle for three months, can lead towards detectable changes in the human brain^16^. A transient increase in gray matter was observed in the occipito-temporal cortex region which comprises the motion sensitive area within the brain. In a more recent study, the effect of mindfulness meditation training in novices identified structural and functional changes in precuneus and posterior default mode network^17^.

In summary, the research to date has shown clear evidences for structural and functional differences in the brain for two healthy groups of subjects considering a particular skill set. Resting-state functional magnetic resonance images (rs-fMRI) could ideally be suited for studying functional differences by helping in understanding **functionally** connected regions when the brain is at rest. This can be combined with magnetic resonance imaging (MRI) to extend functional connections to structural connections to identify local and global changes in the brain. We intend to identify what makes a person proficient at a certain skill or what part of the brain (anatomical/functional) is altered by this skill through analyzing the brain images and further contribute towards classifying two groups of healthy subjects in an expert/amateur paradigm by learning discerning features.

### Summary of Our Contributions

We adapt sparse learning methods (feature reduction and machine learning algorithms) for the brain analysis of different groups of healthy subjects, where one group consists of professional chess players (with many years of experience) and a second group of amateur subjects. Since chess is a demanding board game where cognitive ability is linked to skilled performance; we *hypothesize that there are significant differences, both structurally and in functionally localized brain connections, that could be identified between professional and amateur chess players*. To test this hypothesis, we use brain imaging data curated from grand-master level chess players and controls (amateur chess players)^18^. Our major contributions are the following,

- We analyze morphometric measures on the basis of functional dominance by using a functional parcellation network.
- We propose a novel functional morphometric similarity connectome (FMSC) by combining the anatomical and functional information and enabling sparse learning.
- We classify two groups of healthy controls (chess players) based on their skill specialization (professional and amateur) using the proposed FMSC and hence, identify the structural and functional differences between these groups.

### Related Work

The precuneus was found to be activated during the perception of chess board patterns^19^. To compare anatomical regions in the brain of chess masters and amateur players, a voxel-by-voxel volumetric comparison was performed using a two-sample t-test^20^. A decrease in gray matter volume in the left- and right-caudate regions was found for chess masters compared to amateur controls. Additionally, a resting-state analysis with respect to connections from the caudate also found increased correlations to the posterior cingulate cortex and bilateral angular gyrus. In another study, chess experts demonstrated specific anatomical features in the caudate nucleus, occipito-temporal junction, and the precuneus^21^. A graph theory analysis on rs-fMRI chess data revealed evidence of increased small-world topology and functional connectivity in chess experts compared to amateur players^22^. The functional connectivity matrices were thresholded at different edge strengths to achieve the desired network sparsity. It can be inferred that differences between chess masters and amateur players could exist in the brain topology and functional organization. However, the influence of global topological changes on functional connectivity and vice-versa is not know for such healthy subjects. One way of developing such an understanding would be to study the brain morphology using various non-invasive imaging techniques.

A different approach by Sabuncu et. al.^23^ utilized brain morphology to explain phenotypic variations and identify a global statistical association between brain morphology and observable traits. More recently, the morphometric similarity network (MSN) was proposed to map the network architecture of anatomically connected regions in the brain^24^, where a correlation between morphometric measures and regions of the brain was computed. It was found that MSN modules reiterated known cortical cytoarchitectonic divisions establishing MSN as a valid measure. Structural connectivity can be considered as the basis of functional connectivity and the relationship between these two has been studied in mice^25^, for humans using simulations^26^ and real data^27^, and for specific tasks such as cognition^28^. However, there is a limited evidence of work analyzing the combined effects of anatomical and functional differences. In most cases, anatomical differences were identified and used to localize the search for functional differences.

## Methods

### Data Preparation

#### Anatomical and Functional Imaging Data

The anatomical images were acquired with a repetition time (TR) of 1900 ms, echo time (TE) of 2.26 ms, flip angle of 12°. A total of 176 sagittal slices, each with a slice thickness of 1.0 mm and a voxel size of 1 × 1 × 1 mm, were used. The rs-fMRI images were obtained at TR = 2000 ms, TE = 30 ms, flip angle = 90°. The rs-fMRI data comprised of 205 volumes where 30 whole-head axial slices (from each volume), each 5 mm thick (without gap) and a voxel size of 3.75 × 3.75 × 5 mm, were used. The subjects were instructed to relax with their eyes open and visual fixation on a crosshair centered on the screen. The subject demographics are as follows: the average age and education level (both in years) was 28.67(±9.06) and 13.71(±2.64) for chess masters, respectively and 24.95(±6.14) and 13.87(±2.64) for amateur players, respectively. There were 16 males and 8 females among the chess masters, and 8 males and 15 females among the amateur players. All experiments were carried out by following and conforming to the ethical guidelines and the study was approved by research ethics committee of West China hospital of Sichuan university. An informed consent was taken from all participants involved in the study. More information on the data can be found in the open-source international neuroimaging data-sharing initiative repository^29^.

#### Anatomical data preprocessing

Anatomical features were extracted from high resolution T1-weighted MR images using FreeSurfer^30^. The cortical surface parcellation was performed by following five steps: 1) The MRI volume was registered with the MNI-305 atlas using an affine registration. 2) Bias-field correction and skull stripping were performed. 3) Cutting planes approach was used to remove white- and gray-matter. 4) An initial white surface was generated for each hemisphere which was further refined to follow the intensity gradients between the white- and gray-matter. From this surface, the pial surface was generated by following the intensity gradients between the gray matter and cerebral spinal fluid (CSF). 5) Furthermore, surface labeling was done as in^31^. The parcellation of the cortex for each subject was based on the *17 network functional* parcellation^32^. The metrics of interest, extracted from the cortical parcellations included surface area (SA), gray matter volume (GMV), cortical thickness (CT), curvature index (CI), and folding index (FI). The cortical thickness was measured as the shortest distance from white matter to pial surface. In particular, average cortical thickness (*CT_avg_*) and standard error (*CT_sd_*) was calculated. The rectified mean curvature (MC) was calculated as,

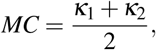

where *κ*_1_, *κ*_2_ are the maximum and minimum curvatures of the surface. The rectified Gaussian curvature (GC) - an intrinsic measure of the curvature of a surface was measured as,

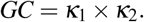

The value of CI represented the maximum intrinsic curvature across all points within the surface. The FI value gives a measure of the local gyrification and was computed as,

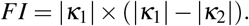

Hence, a total of eight anatomical features were extracted from the T1-weighted MRI data. These metrics have been used in various other studies and shown to be discriminative in general.

#### rs-fMRI data preprocessing

The data preprocessing for rs-fMRI has been intensively explored and discussed by Aurich et. al^33^. In this work, the open-source AFNI (Analysis of Functional NeuroImages) software was used for extraction of functional connectivity networks^34^. The first 5 volumes of the rs-fMRI were discarded to account for the time taken by the tissue to reach steady-state upon application of the magnetic field. Following this, slice timing correction was performed to account for the time difference when each slice was acquired. Motion correction was then performed by aligning all volumes to a reference volume to account for the head movement during the course of the data recording. The functional image was then registered with the corresponding anatomical T1-weighted MR image and smoothing was performed to enable better group-wise analysis. The white matter and ventricle maps from the FreeSurfer output were used as tissue regressors to detect non-BOLD (blood oxygen level dependent) signals in the data. The motion threshold was set to 0.3 mm, with volumes that exceeded this threshold were discarded. Spatial blurring was performed to further reduce the noise and add up coherent signals locally. Finally, band-pass filtering was performed to further isolate the noise with a frequency range of 0.01 – 0.10 Hz.

### Morphometric Similarity Network (MSN)

Morphometric similarity network is a recently proposed method for generating structural connectomes using imaging data from the brain and successfully applied in predicting chronological brain development^24^. The network was generated using diffusion metrics such as fractional anisotropy, mean diffusivity, magnetization transfer and anatomical features including SA, GMV, CT, CI, FI, MC, and GC. For the MSN, the structural connectomes were built using anatomical features. Since correlation requires a feature vector with two or more dimensions, we built several MSNs using combinations of 3 or more anatomical feature vectors. These features have different range of values, e.g., GM and SA values are > 100, whereas GI, FI, and others have values < 10. Therefore each feature (*A*), from a particular region *i*, was *z*-score normalized across all regions as

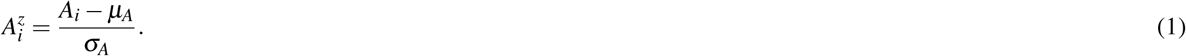

The Pearson’s correlation between feature vectors of different regions (17 networks) was computed to build each MSN. Since we used the Yeo-17 network parcellation map^32^, there were 34 regions of interest across both hemispheres, hence the dimension of the generated MSNs was 34 × 34. The overall network generation process is illustrated in Fig. 1.

**Figure 1.**
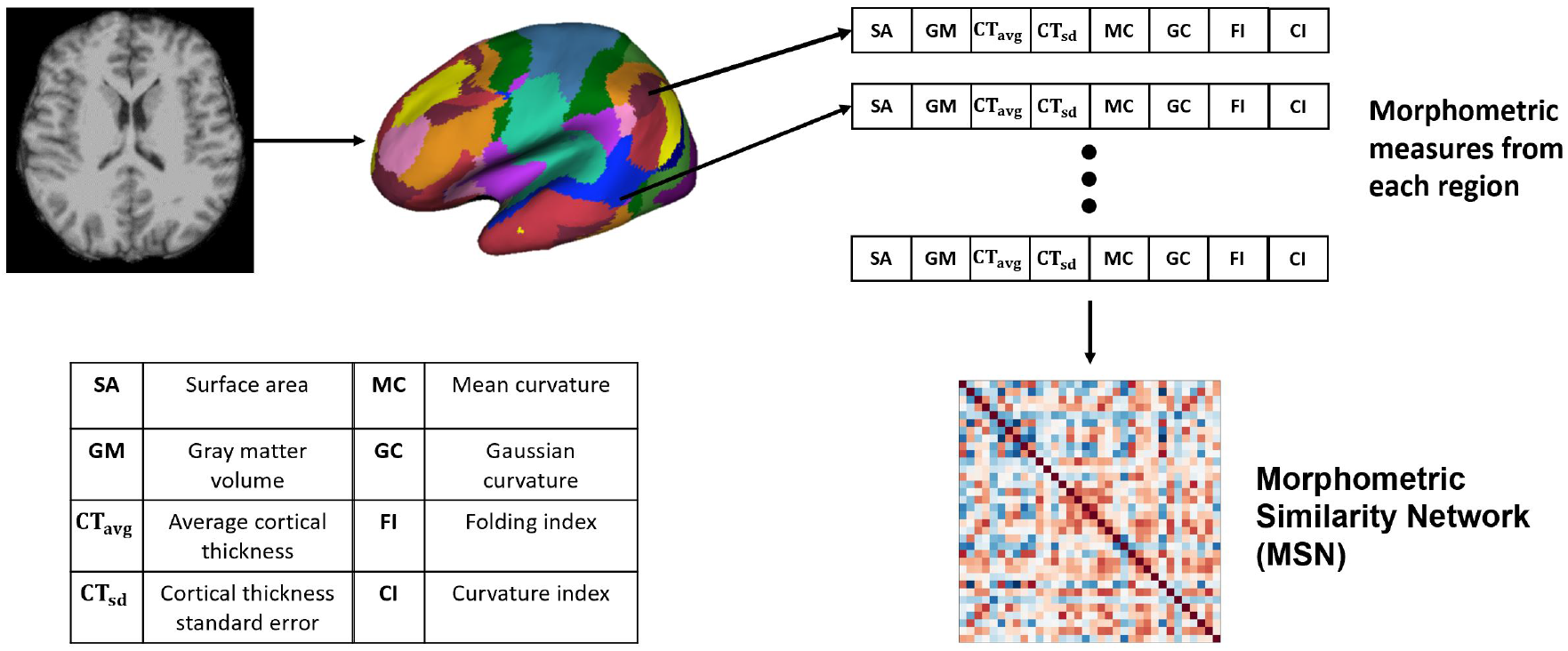
Generating the Morphometric Similarity Network (MSN) from anatomical images where morphometric measures are extracted from the cortical surface of the T1-weighted MRI and Pearson’s correlation is used to build the MSN.

### Functional Connectivity Network (FCN)

Functional connectivity is defined as the correlation of time-series between different voxels or different group of voxels. A FCN can be computed from task-based or rs-fMRI data. In rs-fMRI, no external stimulus is provided while the BOLD signal is recorded to observe the resting-state of the brain. While in task-based evaluation, an external stimulus (finger-tapping, listening to story etc.) is established which alternates with a duration of no stimulus in the experimental design paradigm. For this study, the resting-state paradigm was preferred to examine the general functional differences between the two groups. The Yeo-17 network parcellation^32^ map was overlaid on the preprocessed rs-fMRI images and the average time-series in each functional parcel was extracted. This step amounts to averaging the time-series across all nodes in the parcel. The Pearson’s correlation was then used to compute the correlation between different parcels and generate the functional connectivity network.

### Proposed Functional Morphometric Similarity Connectome (FMSC)

We hypothesize that there are both structural and functional differences between the brains of two distinct healthy groups, and that these differences are highly non-linear and difficult to capture. Hence, we propose a novel approach to combine the anatomical and functional modalities for indirectly building and identifying this relationship (Fig. 2). Towards this, we first extracted morphometric measures: Fig. 2 - Left, and then extracted the FCN. Moreover, since graph metrics enable us to better understand the network topology beyond simple correlation values^35^, therefore we computed the graph metrics - node strength (NS) and node degree (ND), from the FCN. Node degree treats the network as a binary network and represents the number of edges connected to the node which could help in identifying highly connected nodes or hubs in the brain connectome. While node strength, an analog of node degree for weighted networks, represents the sum of the weights of edges connected to the node and helps in identifying the importance of the node.

**Figure 2.**
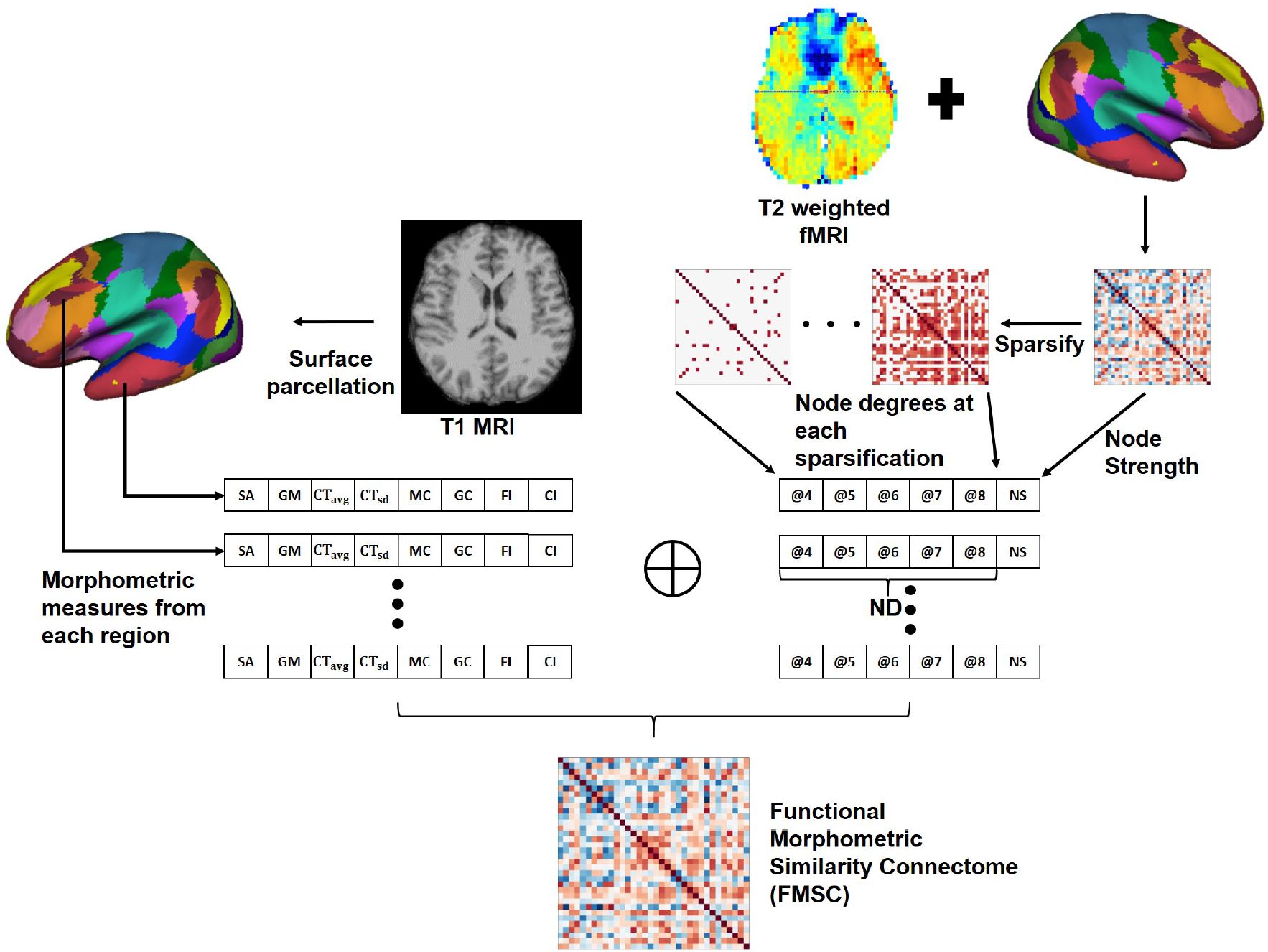
Generating the Functional Morphometric Similarity Connectome (FMSC) from the anatomical and function images. Left: Morphometric measures are extracted from the cortical surface of the T1-weighted MRI. Right: Functional connectivity is generated from the surface of the rs-fMRI and node degrees (ND) are extracted via sparsification (thresholding the edge strength). The morphometric and functional measures are combined to generate the FMSC. NS - node strength, @ - thresholded at.

In particular, we generated sparsified connectivity matrices by thresholding the edge strength of the FCN between [0.4, 0.8], at a step size of 0.1, to successively eliminate weak correlations in the network. This sparsification approach lead to the generation of several undirected connectivity matrices. We then used the ND from these new matrices and the NS from the original FCN as functional measures in place of the Pearson’s correlation (Fig 2 - Right). Finally, we combined the morphometric and functional measures into a single node feature vector which was normalized using the *z* – *score* (Eq 1). The correlation between different nodes was computed using this feature vector to build the novel connectivity matrix, the functional morphometric similarity connectome.

## Results

### Statistical analysis

To identify whether morphometric measures (derived from functionally defined anatomical regions) are distinct between chess masters and amateur players, we performed a two-sample t-test for each brain region and metric (See Figure 1). The significance value was set at p < 0.05, and a family-wise error (FWE) correction based on the Holm-Sidak method^36^ was employed to control the number of Type-I errors, while also reducing the increased risk (due to correction) of Type-II errors. Two significant regions were identified along with cortical thickness as the metric of interest. Table 1 shows the regions and the FWE corrected p-values. It should be noted that these regions (somatomotor A and peripheral visual) are different from those previously identified in literature, which shows the benefit of using functional parcellation. Since the chosen parcellation comprises of spatially non-contiguous brain regions, we performed the t-test with the anatomical parcellation atlas - Destrieux atlas^37^, which has 74 gyral and sulcal regions. Among these regions, six regions were found statistically significantly different between groups and are shown in Table 1. These six regions are known to be part of the somatosensory (pre-central gyrus, central sulcus, and supra-marginal gyrus) and central execution (gyrus rectus) networks. This is in agreement with neuroscience studies that have shown that highly complex games, such as chess, favors the development of the frontal lobe^38^.

**Table 1.**
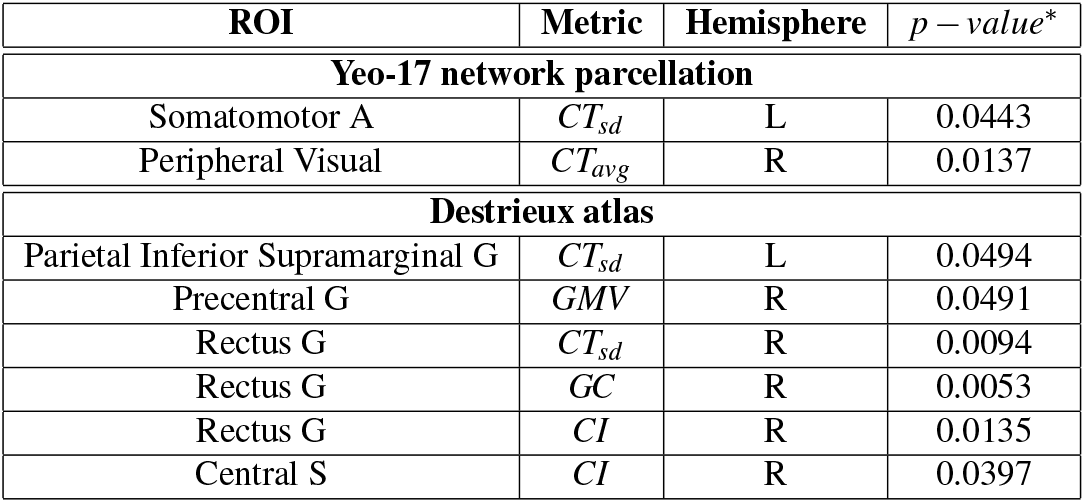
Anatomically significant functionally defined regions and the corresponding metric of difference. * - FWE corrected, L - Left, R - Right, G - Gyrus, S - Sulcus, *CT_sd_* - standard error of cortical thickness, *CT_avg_* - Average cortical thickness, *GMV* - Gray volume, *GC* - Gaussian curvature, *CI* - Curvature Index, ROI - region of interest.

| ROI | Metric | Hemisphere | $p$ - value* |
| --- | --- | --- | --- |
| **Yeo-17 network parcellation** |  |  |  |
| Somatomotor A | $CT_{sd}$ | L | 0.0443 |
| Peripheral Visual | $CT_{avg}$ | R | 0.0137 |
| **Destrieux atlas** |  |  |  |
| Parietal Inferior Supramarginal G | $CT_{sd}$ | L | 0.0494 |
| Precentral G | $GMV$ | R | 0.0491 |
| Rectus G | $CT_{sd}$ | R | 0.0094 |
| Rectus G | $GC$ | R | 0.0053 |
| Rectus G | $CI$ | R | 0.0135 |
| Central S | $CI$ | R | 0.0397 |

### Classification

After identifying statistically significantly different regions, we generated the connectivity matrix to identify further differences between two healthy groups of individuals. To ensure a strong classification model, the *k*–fold cross-validation approach^33^ was used with the value of *k* = 10. Note that cross-validation is an approach to do out-of-sample testing with limited data wherein the data is split into training and testing sets to enable model validation. In *k*– fold cross-validation, the data is split into *k* different training and test sets and the average performance across all metrics is the overall model evaluation. We used a stratified 10-fold cross-validation technique, where the data splitting ensures that there are a similar number of samples from all classes in the test set.

All connectivity matrices/connectomes generated were symmetrical i.e., they were identical across the diagonal. The diagonal itself represented self-correlation and was always equal to 1. Thus, only the upper-right triangle was extracted from these connectivity matrices and the resulting feature vector was used in the classification task. For an *m* × *m* matrix the feature vector dimension was computed as 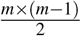. Since the networks were 34 × 34, the extracted feature vector was of dimension 561. The feature dimension was greater than the size of the training data and thus, feature selection was used to reduce the feature vector size. Towards this, a simple student’s t-test was performed and only significant features (at a confidence interval of *p* < 0.05) were chosen. The selected features were then used to train a support vector machine (SVM)^39^ with a linear kernel, regularization and scaling parameters were set to *C* = 1 and *γ* = 1*e* – 4, respectively. Note that any other off-the-shelf classifier can be used for this purpose, however, the classification performance of SVM is at par. The performance was evaluated using accuracy, precision, and recall. These evaluation metrics are commonly used and widely known in the literature; however, to make the manuscript self-contained, we briefly summarize them as follows.

Accuracy is defined as the ratio of the total number of correctly classified items to the total number of items and was computed as

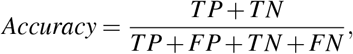

where TP - true positives, FP - false positives, TN - true negatives and FN - false negatives. Precision is defined as the ratio of correctly predicted positive results to the total predicted positive results and was computed as

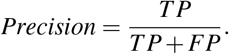

F1-score is defined as the harmonic mean of precision and recall and was computed as

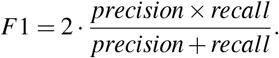

#### FCN-based classification

The effect of feature selection on FCN-based classification was evaluated by first performing a 10-fold cross-validation using all the extracted features. As a result, SVM was trained using 561 features, resulting in the classification performance presented in Table 2. Later, feature selection (via t-test) was performed prior to training the SVM in each fold. This resulted in an average accuracy of 76.33%, a significant improvement (of over 11%) compared to the baseline. The common significant connections across all 10-folds of the cross-validation were identified and are shown in Fig. 3a. Lateral and medial views show the intra-hemispheric connections and inter-hemispheric connections are presented in the dorsal view.

**Table 2.** 10-fold cross-validation performance using functional connectivity features. The bold values show that the performance improved in all matrices using feature selection. w/o - without, w - with.

| Approach | Accuracy (%) | Precision (%) | F1-score (%) |
| --- | --- | --- | --- |
| w/o t-test | 64.83 | 62.67 | 64.83 |
| **w t-test** | **76.33** | **76.83** | **74.05** |

**Figure 3.**
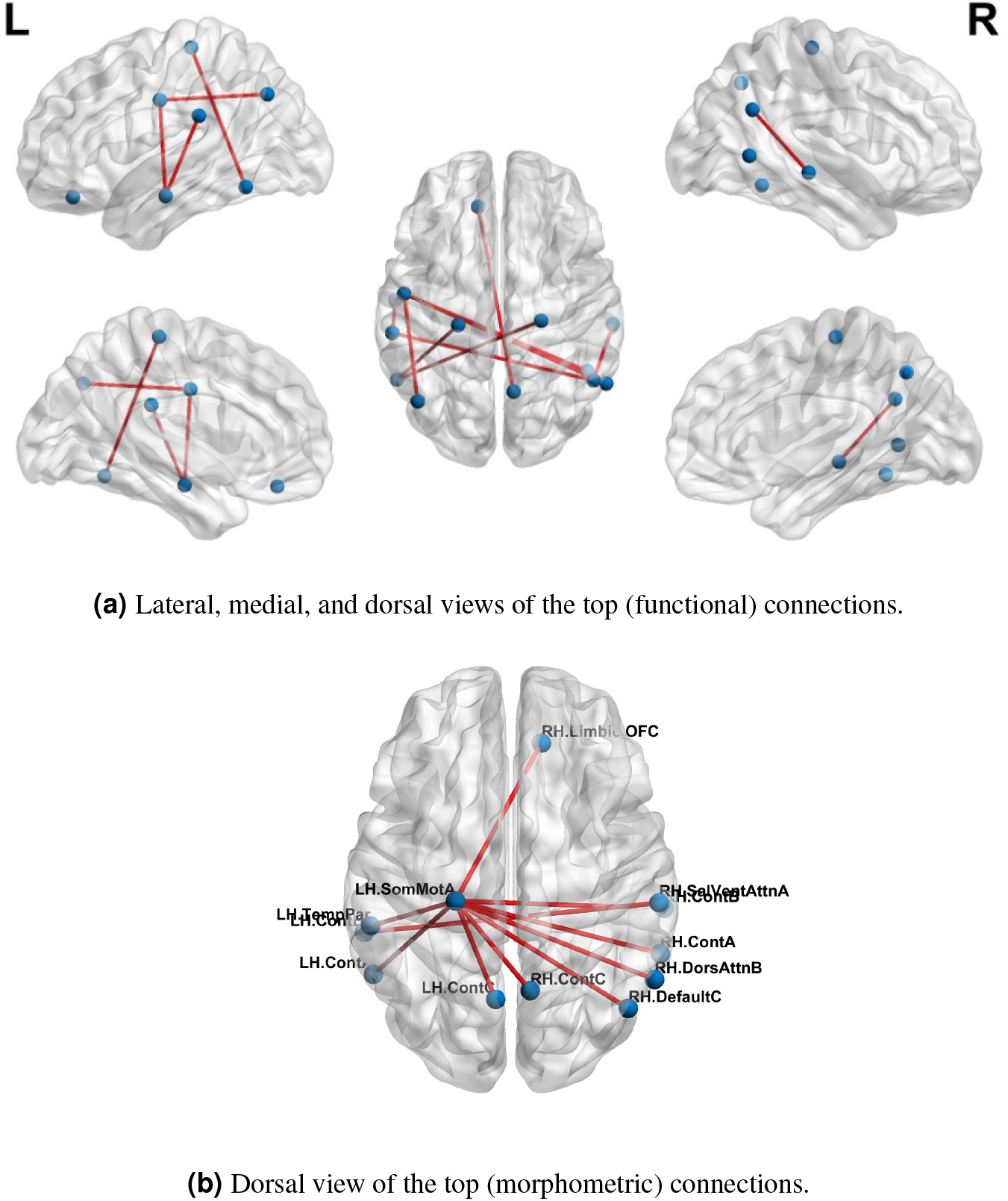
Top-10 connections identified in classification of chess masters vs amateur players. (a) Functional connections (b) Morphometric connections. Table 4 lists the names of the ROIs in these top connections.

#### MSN-based classification

Having established the benefit of using feature selection towards the classification task and our baseline classification accuracy, we performed classification using connectivity features from the MSN networks. Since we consider the construction of MSNs using 3 or more measures, the total number of MSNs was computed as

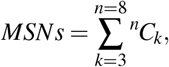

where the total number of morphometric measures was 8. There were a total of 219 MSNs and thus, 219 trained SVM models. Among these models, the best 10-fold classification accuracy was achieved using SA, CT_sd_ and CI as the morphometric features of interest. The MSN representing the connections for this particular model is shown in Fig. 3b. We found that the saliency and ventral attention network in the left hemisphere and the dorsal attention network in the right hemisphere had significantly different connections across networks.

#### FMSC-based classification

We validated our hypothesis that MSN- and FCN-based approaches contribute complementary information using a simple majority voting strategy, called pseudo-FMSC, combining the best performing MSN- and FCN-based classification models. The common connections across the majority voting model are shown in Fig. 4a. The saliency and ventral attention network, dorsal attention network, and central visual network were common across the models. The resulting model had a 6% and 3% improvement in classification performance over the best performing MSN and FCN models, respectively, reaching an overall accuracy of 80%. This indicated the presence of potentially complementary information in the MSN to that of FCN. Thus, we combined the MSN and FCN metrics via feature concatenation. The FCN was thresholded at edge strengths of [0.4, 0.5, 0.6, 0.7, 0.8] and NDs were computed for each of these sparse un-directed networks, whereas NS was computed directly from the FCN. The total number of features was now reduced to 14 (8 morphometric measures and 6 functional measures). Similar to MSN-based classification, we tested all possible combinations and this resulted in 364 (^14^*C*_3_) different models. The highest accuracy of 88% was achieved and Fig. 4b shows the common significant connections across the 10-fold cross-validation in the best performing FMSC model.

**Figure 4.**
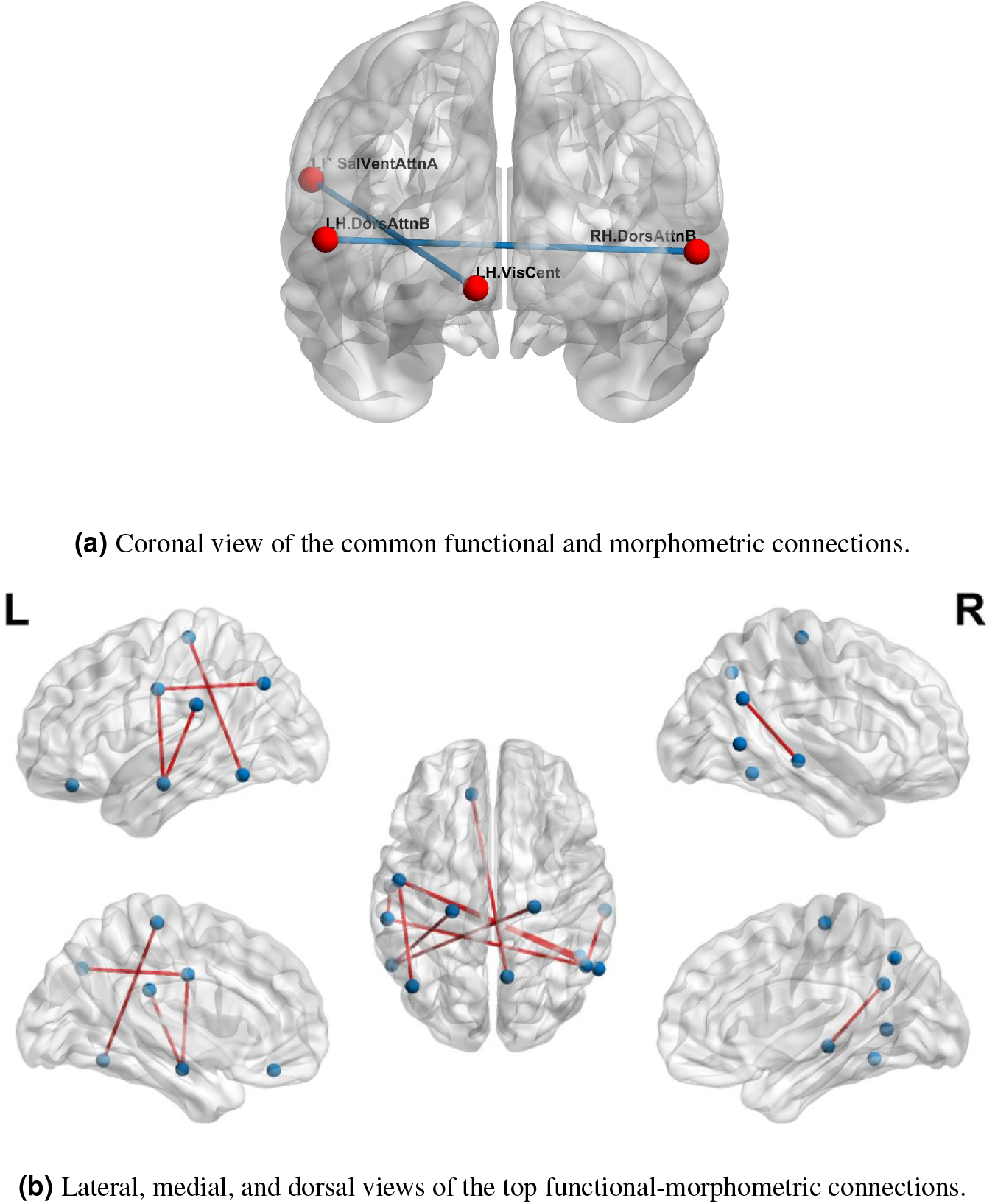
Functional anatomical connections differentiating chess masters from amateur players. (a) Common functional and morphometric connections (b) Functional-morphometric connections. Table 4 lists the names of the ROIs in these top connections.

## Discussion

In this work, we have proposed a novel approach (FMSC) for combining metrics from anatomical and functional brain images, to identify distinct connectomes between two groups of healthy subjects, specifically chess masters and amateur players. Our trained machine learning-based model achieved a classification accuracy of 88% showing drastic improvements over the standard functional connectivity-based approach. The performance measures for the top performing models are presented in Table 3. It should be noted that the best model (using sparsification-based FMSC) significantly improves the performance (accuracy) by ≈ 14% when compared with MSN-based approach, and ≈ 11% when compared with FCN-based approach. These results add credence to the success of the proposed FMSC-based classification.

**Table 3.**
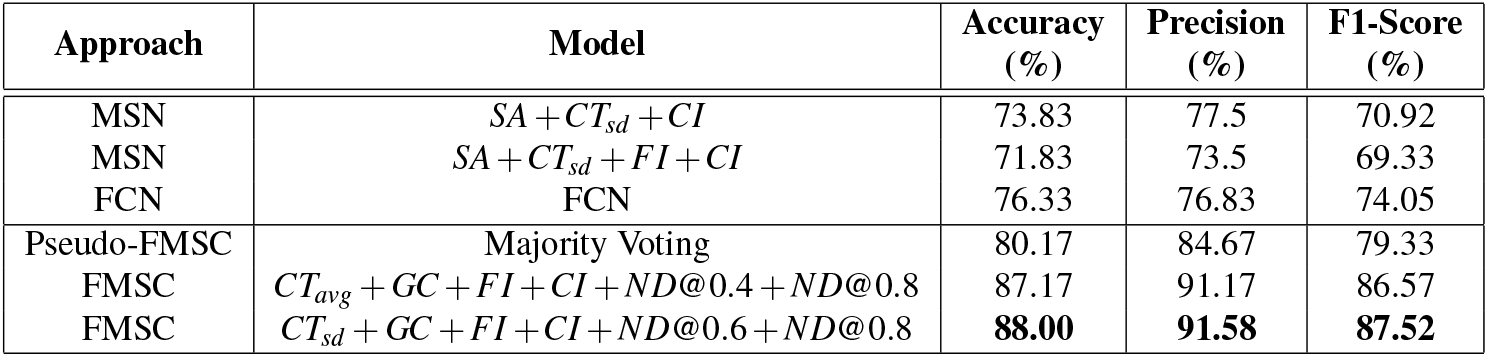
Classification results of morphometric similarity networks (MSN), functional connectivity networks (FCN) and functional morphometric similarity (FMSC) on 10-fold cross-validation. SA - Surface area, CT_avg - Average cortical thickness, CT_sd - Cortical thickness standard error, FI - Folding index, CI - Curvature Index, GC - Gaussian curvature, ND@ - Node degree at threshold level.

| Approach | Model | Accuracy (%) | Precision (%) | F1-Score (%) |
| --- | --- | --- | --- | --- |
| MSN | $SA + CT_{sd} + CI$ | 73.83 | 77.5 | 70.92 |
| MSN | $SA + CT_{sd} + FI + CI$ | 71.83 | 73.5 | 69.33 |
| FCN | FCN | 76.33 | 76.83 | 74.05 |
| Pseudo-FMSC | Majority Voting | 80.17 | 84.67 | 79.33 |
| FMSC | $CT_{avg} + GC + FI + CI + ND@0.4 + ND@0.8$ | 87.17 | 91.17 | 86.57 |
| FMSC | $CT_{sd} + GC + FI + CI + ND@0.6 + ND@0.8$ | **88.00** | **91.58** | **87.52** |

Additionally, we have used functional parcellation that combines functionally coupled regions into the same networks. This allowed us to study whether broadly defined functional networks show morphometric differences. We identified two such coupled regions with statistically significant differences in cortical thickness. The network analysis using the functional parcellation enabled us to look at higher-order connectivity across anatomical regions which are non-local, owing to the functionally defined parcellation. We analyzed the use of morphometric similarity networks towards the classification and found an accuracy above that of a simple coin flip. This further indicates that there are anatomical differences between non-local regions and that the relationship may not be a simple linear relationship.

The proposed FMSC-based classification shows that there exists a complex functional-morphometric relationship. To establish whether functional-morphometric measures provide a unique look into the organization, we examined the overlap of top connections between the best MSN, FCN, and FMSC models. The metrics used in each of these networks were different; therefore; we expected very few regions to be common. We identified an atypical within- and between-hemisphere dorsal attention network (DAN) connection. It should be noted that DAN is a task-positive network that is cued during externally directed attention tasks^40^. We also studied the top-10 connections in the MSN-, FCN-, and FMSC-based classification and are presented in Table 4. The saliency and ventral attention (SVA) network was identified to be in the top-10 connections in both the FCN and FMSC-based approaches, serving as a kind of hub to connections that were significantly different between chess masters and amateur players. The SVA network commonly serves as a switch between the default mode network and the central execution network. The between-hemisphere connection of the SVA and the control network was identified to be functionally different. We also found that the right hemisphere control network is common in both FCN-based and FMSC-based networks as a kind of hub. A minimal overlap between the significant connections across different approaches validates the hypothesis that there are indirect and non-linear relationships between anatomy and function. This indirect relationship can be identified using the proposed FMSC and potentially help in an early diagnosis of several neurological disorders.

**Table 4.**
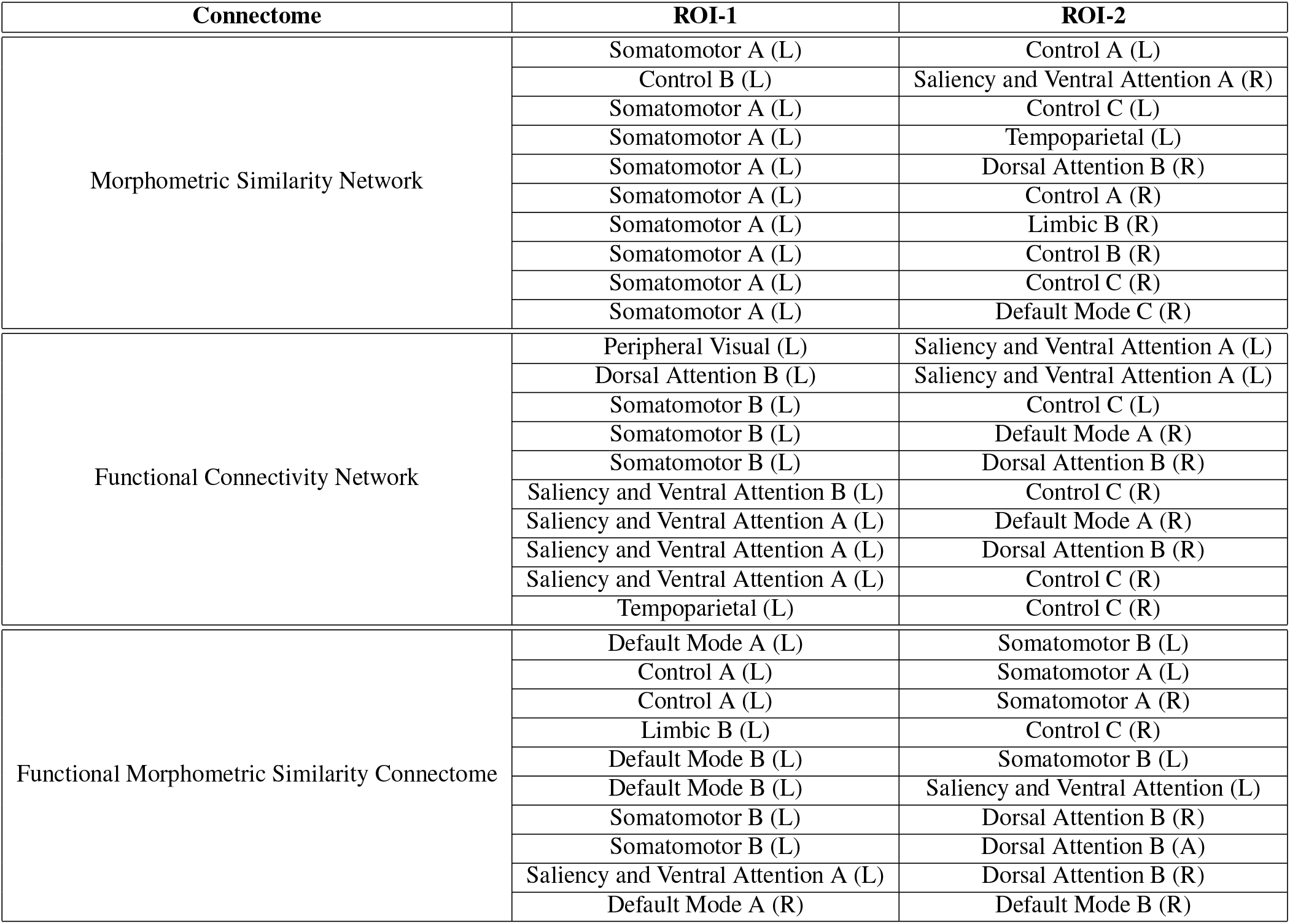
Top-10 significant connections based on morphometery, functionality, and functional-morphometery. ROI - Region of Interest, L - Left, R - Right.

| Connectome | ROI-1 | ROI-2 |
| --- | --- | --- |
| Morphometric Similarity Network | Somatomotor A (L) | Control A (L) |
|  | Control B (L) | Saliency and Ventral Attention A (R) |
|  | Somatomotor A (L) | Control C (L) |
|  | Somatomotor A (L) | Tempoparietal (L) |
|  | Somatomotor A (L) | Dorsal Attention B (R) |
|  | Somatomotor A (L) | Control A (R) |
|  | Somatomotor A (L) | Limbic B (R) |
|  | Somatomotor A (L) | Control B (R) |
|  | Somatomotor A (L) | Control C (R) |
|  | Somatomotor A (L) | Default Mode C (R) |
| Functional Connectivity Network | Peripheral Visual (L) | Saliency and Ventral Attention A (L) |
|  | Dorsal Attention B (L) | Saliency and Ventral Attention A (L) |
|  | Somatomotor B (L) | Control C (L) |
|  | Somatomotor B (L) | Default Mode A (R) |
|  | Somatomotor B (L) | Dorsal Attention B (R) |
|  | Saliency and Ventral Attention B (L) | Control C (R) |
|  | Saliency and Ventral Attention A (L) | Default Mode A (R) |
|  | Saliency and Ventral Attention A (L) | Dorsal Attention B (R) |
|  | Saliency and Ventral Attention A (L) | Control C (R) |
|  | Tempoparietal (L) | Control C (R) |
| Functional Morphometric Similarity Connectome | Default Mode A (L) | Somatomotor B (L) |
|  | Control A (L) | Somatomotor A (L) |
|  | Control A (L) | Somatomotor A (R) |
|  | Limbic B (L) | Control C (R) |
|  | Default Mode B (L) | Somatomotor B (L) |
|  | Default Mode B (L) | Saliency and Ventral Attention (L) |
|  | Somatomotor B (L) | Dorsal Attention B (R) |
|  | Somatomotor B (L) | Dorsal Attention B (A) |
|  | Saliency and Ventral Attention A (L) | Dorsal Attention B (R) |
|  | Default Mode A (R) | Default Mode B (R) |

In future work, we will use a further splitting of the network labels to analyze morphometric features of individual regions within the functional cluster. Moreover, with the increased dimensionality of the feature vectors, deep learning algorithms can be utilized to perform classification but the size of the dataset may prove to be a limiting factor^41^. While MSN enabled us to combine structural information, as a potential extension of this work, we would combine structural information from diffusion tensor imaging based measures such as fractional anisotropy, mean diffusivity, and structural node degree into the morphometric measures to further strengthen the functional-morphometric relationship.

## Author contributions statement

All authors reviewed the manuscript and agreed on the final version.

## References

1. Zuo, X.-N. & Xing, X.-X. Test-retest reliabilities of resting-state fmri measurements in human brain functional connectomics: a systems neuroscience perspective. Neurosci. & Biobehav. Rev. 45, 100–118 (2014).

2. Dosenbach, N. U. et al. Prediction of individual brain maturity using fmri. Science 329, 1358–1361 (2010).

3. Thompson, P. & Apostolova, L. Computational anatomical methods as applied to ageing and dementia. The Br. journal radiology (2014).

4. Chen, G., Saad, Z. S., Britton, J. C., Pine, D. S. & Cox, R. W. Linear mixed-effects modeling approach to fmri group analysis. Neuroimage 73, 176–190 (2013).

5. Wee, C.-Y. et al. Identification of mci individuals using structural and functional connectivity networks. Neuroimage 59, 2045–2056 (2012).

6. Chen, X. et al. High-order resting-state functional connectivity network for mci classification. Hum. brain mapping 37, 3282–3296 (2016).

7. He, T. et al. Is deep learning better than kernel regression for functional connectivity prediction of fluid intelligence? In 2018 International Workshop on Pattern Recognition in Neuroimaging (PRNI), 1–4 (IEEE, 2018).

8. Casanova, R. et al. High dimensional classification of structural mri alzheimer’s disease data based on large scale regularization. Front. neuroinformatics 5, 22 (2011).

9. Wang, Y. et al. A novel approach for stable selection of informative redundant features from high dimensional fmri data. arXiv preprint arXiv:1506.08301 (2015).

10. Munsell, B. C. et al. Evaluation of machine learning algorithms for treatment outcome prediction in patients with epilepsy based on structural connectome data. NeuroImage 118, 219–230 (2015).

11. Basten, U., Hilger, K. & Fiebach, C. J. Where smart brains are different: A quantitative meta-analysis of functional and structural brain imaging studies on intelligence. Intelligence 51, 10–27 (2015).

12. Rioult-Pedotti, M.-S., Friedman, D., Hess, G. & Donoghue, J. P. Strengthening of horizontal cortical connections following skill learning. Nat. neuroscience 1, 230–234 (1998).

13. Gaser, C. & Schlaug, G. Brain structures differ between musicians and non-musicians. J. Neurosci. 23, 9240–9245 (2003).

14. Maguire, E. A., Woollett, K. & Spiers, H. J. London taxi drivers and bus drivers: a structural mri and neuropsychological analysis. Hippocampus 16, 1091–1101 (2006).

15. Di, X. et al. Altered resting brain function and structure in professional badminton players. Brain connectivity 2, 225–233 (2012).

16. Driemeyer, J., Boyke, J., Gaser, C., Büchel, C. & May, A. Changes in gray matter induced by learning—revisited. PloS one 3 (2008).

17. Yang, C.-C. et al. Alterations in brain structure and amplitude of low-frequency after 8 weeks of mindfulness meditation training in meditation-naïve subjects. Sci. reports 9, 1–10 (2019).

18. Li, K. et al. A multimodal mri dataset of professional chess players. Sci. data 2, 150044 (2015).

19. Wan, X. et al. The neural basis of intuitive best next-move generation in board game experts. Science 331, 341–346 (2011).

20. Duan, X. et al. Reduced caudate volume and enhanced striatal-dmn integration in chess experts. Neuroimage 60, 1280–1286 (2012).

21. Hänggi, J., Brütsch, K., Siegel, A. M. & Jäncke, L. The architecture of the chess player’s brain. Neuropsychologia 62, 152–162 (2014).

22. Duan, X. et al. Functional organization of intrinsic connectivity networks in chinese-chess experts. Brain research 1558, 33–43 (2014).

23. Sabuncu, M. R. et al. Morphometricity as a measure of the neuroanatomical signature of a trait. Proc. Natl. Acad. Sci. 113, E5749–E5756 (2016).

24. Seidlitz, J. et al. Morphometric similarity networks detect microscale cortical organization and predict inter-individual cognitive variation. Neuron 97, 231–247 (2018).

25. Grandjean, J., Zerbi, V., Balsters, J. H., Wenderoth, N. & Rudin, M. Structural basis of large-scale functional connectivity in the mouse. J. Neurosci. 37, 8092–8101 (2017).

26. Stam, C. et al. The relation between structural and functional connectivity patterns in complex brain networks. Int. J. Psychophysiol. 103, 149–160 (2016).

27. Uddin, L. Q. Complex relationships between structural and functional brain connectivity. Trends cognitive sciences 17, 600–602 (2013).

28. Zimmermann, J., Griffiths, J. D. & McIntosh, A. R. Unique mapping of structural and functional connectivity on cognition. J. Neurosci. 38, 9658–9667 (2018).

29. INDI. A multimodal mri dataset of professional chess players. (2015). (last accessed 2020-06-06).

30. Fischl, B. Freesurfer. Neuroimage 62, 774–781 (2012).

31. Desikan, R. S. et al. An automated labeling system for subdividing the human cerebral cortex on mri scans into gyral based regions of interest. Neuroimage 31, 968–980 (2006).

32. Thomas Yeo, B. et al. The organization of the human cerebral cortex estimated by intrinsic functional connectivity. J. neurophysiology 106, 1125–1165 (2011).

33. Arlot, S., Celisse, A. et al. A survey of cross-validation procedures for model selection. Stat. surveys 4, 40–79 (2010).

34. Cox, R. W. Afni: software for analysis and visualization of functional magnetic resonance neuroimages. Comput. Biomed. research 29, 162–173 (1996).

35. Bullmore, E. & Sporns, O. Complex brain networks: graph theoretical analysis of structural and functional systems. Nat. reviews neuroscience 10, 186–198 (2009).

36. Šidák, Z. Rectangular confidence regions for the means of multivariate normal distributions. J. Am. Stat. Assoc. 62, 626–633 (1967).

37. Destrieux, C., Fischl, B., Dale, A. & Halgren, E. Automatic parcellation of human cortical gyri and sulci using standard anatomical nomenclature. Neuroimage 53, 1–15 (2010).

38. Ortiz-Pulido, R., Ortiz-Pulido, R., García-Hernández, L. I., Pérez-Estudillo, C. A. & Ramírez-Ortega, M. L. Neuroscientific evidence support that chess improves academic performance in school. Revista Mexicana de Neurociencia 20, 194–199 (2019).

39. Cortes, C. & Vapnik, V. Support-vector networks. Mach. learning 20, 273–297 (1995).

40. Spreng, R. N., Stevens, W. D., Chamberlain, J. P., Gilmore, A. W. & Schacter, D. L. Default network activity, coupled with the frontoparietal control network, supports goal-directed cognition. Neuroimage 53, 303–317 (2010).

41. RaviPrakash, H., Watane, A., Jambawalikar, S. & Bagci, U. Deep learning for functional brain connectivity: Are we there yet? In Deep Learning and Convolutional Neural Networks for Medical Imaging and Clinical Informatics, 347–365 (Springer, 2019).

